# Sex-Dependent Cross-Resilience to Social Defeat and Learned Helplessness: The Role of BDNF-ERK Signaling and Norepinephrine

**DOI:** 10.1101/2025.10.27.684864

**Authors:** Maryia Bairachnaya, Lucy Penney, Gohar Fakhfouri, Elsa Isingrini, Bruno Giros

## Abstract

Major depressive disorder (MDD) and post-traumatic stress disorder (PTSD) are two of the most prevalent and disabling psychiatric conditions, both closely tied to stress exposure. While they have distinct diagnostic criteria, MDD and PTSD share considerable comorbidity, overlapping behavioral symptoms, and common neurobiological pathways. However, not all individuals exposed to chronic or traumatic stress develop these disorders, highlighting the importance of resilience, the capacity to maintain psychological and physiological stability under stress. An emerging yet understudied concept is cross-resilience: the idea that resilience to one form of stress may confer protection against another. While animal models have effectively captured individual differences in stress responses, few have explored whether resilience generalizes across distinct stress modalities or differs by sex.

We examined cross-resilience using two validated rodent models: chronic social defeat stress (CSDS) and learned helplessness (LH). These paradigms model complementary aspects of stress vulnerability. We assessed how prior CSDS experience influenced subsequent responses to LH. We also evaluated the role of the noradrenergic system using wild-type and norepinephrine (NE)-deficient (VMAT2^loxDBHcre^ KO) mice. Behavioral phenotyping was combined with molecular analyses in key brain regions involved in emotion, motivation, and fear processing: the ventral tegmental area (VTA), nucleus accumbens (NAc), and amygdala (AMY). We focused on BDNF and ERK/MAPK signaling pathways, known to mediate neuroplasticity and stress resilience.

Our findings reveal sex-specific patterns of cross-resilience, with prior CSDS resilience predicting LH resilience in males but not females. Molecular results indicate distinct adaptations across brain regions and sexes, underscoring the biological complexity of resilience. These insights may inform personalized strategies for preventing and treating stress-related disorders.

## Introduction

Major depressive disorder (MDD) and post-traumatic stress disorder (PTSD) are among the most prevalent and debilitating psychiatric conditions. Although they have distinct diagnostic criteria, both are strongly linked to stress exposure as a primary risk factor and share common disruptions at molecular, cellular, and circuit levels. Globally, MDD affects 5-20% of the population (Smith, 2014), and PTSD affects approximately 4-5% (White et al., 2015; Koenen et al., 2017). This suggests that not all individuals exposed to highly stressful or traumatic events develop psychopathology (Rutter, 2006). Such variability is commonly attributed to individual **resilience to stress** – the capacity to adapt successfully to adversity and mitigate its negative consequences. Resilience is not simply the absence of pathology, but a dynamic process involving active molecular, hormonal, cellular, and behavioral mechanisms that maintain or restore homeostasis after stress exposure (Russo et al., 2012)

Approximately half of individuals with PTSD also experience MDD (Kessler et al., 1995; Breslau et al., 2000; Rytwinski et al., 2013; Flory and Yehuda, 2015), indicating a high rate of comorbidity between the two disorders. This comorbidity is associated with poorer treatment outcomes (Green et al., 2006; Galovski et al., 2016) and an increased risk of suicidal behavior (Oquendo et al., 2003; Kessler et al., 2005). These findings suggest that MDD and PTSD may share overlapping mechanisms of vulnerability and resilience. Understanding these shared mechanisms is crucial, as there are currently no targeted interventions for individuals affected by both disorders (Flory and Yehuda, 2015).

One phenomenon that remains poorly understood is **cross-resilience**. It refers to the idea that resilience to one type of stress may confer protection against another. While animal studies have extensively examined resilience in response to either chronic or trauma-related stress, few have investigated resilience to sequential stressors within the same individuals, and even fewer have considered sex differences in these responses. This study addresses this gap by investigating cross-resilience between two well-established and translationally relevant animal models: **chronic social defeat stress (CSDS)** and **learned helplessness (LH)**. Both models are well validated for inducing behavioral, molecular, and circuit-level changes that mirror key aspects of human stress-related disorders (van der Kolk et al., 1985; Chourbaji et al., 2005; Hollis and Kabbaj, 2014; Silveira and Joca, 2023). Using these paradigms, we aim to examine how exposure to one form of stress may influence susceptibility or resilience to another and to uncover the molecular mechanisms underlying this interaction.

Given the pivotal role of the noradrenergic (NE) system in orchestrating stress responses and promoting resilience, we focus on its contribution to cross-resilience. NE signaling has been implicated in adaptive stress responses in both humans (Gilam et al., 2017; Naegeli et al., 2018; Grueschow et al., 2021) and animal models (Isingrini et al., 2016; Kim et al., 2016; Zhang et al., 2019; Zhai et al., 2023). Moreover, NE modulates neuronal plasticity through mechanisms such as the upregulation of brain-derived neurotrophic factor (BDNF) (Juric et al., 2008; Yaniv et al., 2010) and the activation of mitogen-activated protein kinase (MAPK/ERK) pathways (Alblas et al., 1993; Tolbert et al., 2003; Sun et al., 2024), both of which are critical for synaptic plasticity and resilience. Dysregulation of ERK and BDNF signaling has been linked to the pathophysiology of MDD (Dwivedi et al., 2001; Duric et al., 2010; Malki et al., 2015) and PTSD (Xiao et al., 2011; Zhang et al., 2014; Zhao et al., 2020; Chaaya et al., 2021), and BDNF expression has been shown to differ between resilient and susceptible mice following CSDS (Mallei et al., 2019).

To investigate the role of NE signaling in cross-resilience-related plasticity, we employ a genetically engineered mouse model with selective knockout of NE transmission (VMAT2^loxDBHcre^ KO aka NEKO), previously shown to display increased susceptibility to CSDS in males (Isingrini et al., 2016).

Altogether, we compare male and female mice as well as wild-type and NE-deficient mice, to examine molecular signaling cascades in key brain regions involved in stress regulation, motivation, and emotional processing. These include the ventral tegmental area (VTA) and nucleus Accumbens (NAc) critical for reward and the amygdala (AMY), which is essential for emotional and fear-related responses. These regions receive dense NE innervation (Sara, 2009) and exhibit stress-induced changes in ERK and BDNF activity (Iñiguez et al., 2010; Lakshminarasimhan and Chattarji, 2012; Maldonado et al., 2014; Wook Koo et al., 2016; Pagliusi et al., 2018; Contesse et al., 2021; Pagliusi et al., 2022; Bastos et al., 2024).

By elucidating the sex- and NE-dependent mechanisms of cross-resilience, this study seeks to uncover new insights into the biological processes that mediate stress adaptation and contribute to the onset - or prevention - of MDD and PTSD.

## Methods

### Animals

C57BL/6 male and female mice (8-11 weeks old) and CD-1 retired breeder adult males (3-month-old) were obtained from Charles River laboratories. The floxed *VMAT2* mouse strain generated at the Mouse Clinical Institute (Institut Clinique de la Souris, MCI/ICS, Illkirch, France) was breed in our animal facility; the DBHcre mouse strain (B6.FVB(Cg)-*Tg(Dbh-cre)KH212Gsat*/Mmucd; stock number 031028-UCD) was supplied by the Mutant Mouse Regional Resource Centers (MMRRC). The brain-specific NE-depleted mice (*VMAT2^DBHcre^KO*) were generated by crossing heterozygous *VMAT2* floxed mice (VMAT2^lox/+^) with heterozygous DBH^cre/+^ mice to obtain double heterozygous offspring which then was crossed to generate the mice used for the behavioral experiments (only KO and WT mice were used).

The colonies were housed in the animal facility of the Douglas Research Centre. The mice were kept under standard conditions at 22±1°C, 60% relative humidity, and a 12-hour light/dark cycle with food and water available *ad libitum*.

All procedures were done in accordance with the Canadian Council on Animal Care guidelines (CCAC; https://ccac.ca/en/guidelines-and-policies/the-guidelines/) and approved by the Animal Care Committee of the Douglas Research Centre.

### Chronic Social Defeat Stress

#### Males

The test was performed as previously described (Golden et al., 2011). Prior to beginning the defeats, CD-1 aggressor mice were screened over three consecutive days and selected according to their latency (under 60 sec) and consistency to attack (two consecutive screening days) a novel C57BL/6 male mouse placed in their home cage.

Experimental C57BL/6 male mice were subjected to a daily 5-minutes defeat with a different CD-1 mouse on each of 10 consecutive days (**Fig. 1B**). A defeat lasted for either 5 minutes or 10 attacks, whichever occurred first. The defeats were carried out in large rat cages with two halves separated by a plexiglass divider. After each defeat, the mouse remained in the aggressor’s home cage, but separated from the aggressor by a perforated translucent Plexiglas divider, allowing the defeated mouse to have visual, olfactory and auditory interaction with the aggressor until the next defeat. Control mice were housed in pairs separated by a Plexiglas divider and were moved to a different cage each day, but never had physical contact with each other and they were not exposed to CD-1 aggressors. The controls were housed in a different room to the defeated mice, where there were no CD-1 mice present.

**Figure 1.**
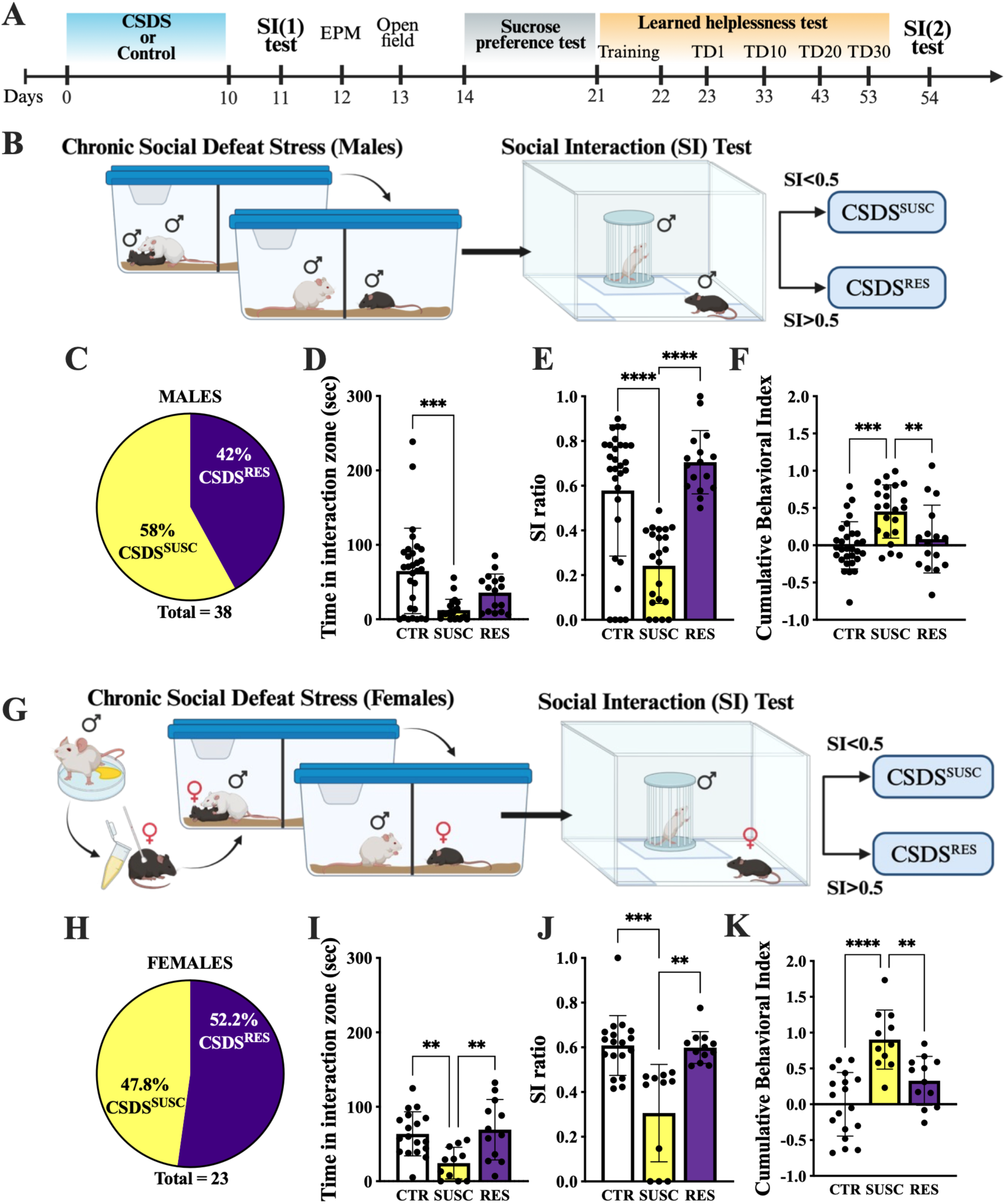
Behavior of male and female stressed and control groups in social interaction test. A. Experimental timeline for studying cross-resilience for chronic social defeat stress (CSDS) and learned helplessness (LH). Graphic representation of CSDS paradigm for males (B) and females (G). The proportion of CSDS resilient and susceptible animals for males (C) and females (H). The time spent in the interaction zone when the target mouse was present for males (D) and females (I). Social interaction ratio for males (E) and females (J). Cumulative behavioral index for males (F) and females (K). Data are shown as mean ± SD, with \**p*<0.05, \*\**p*<0.01, ***p<0.001, and ****p<0.0001 indicating significance. The images A, B and G were created in BioRender.

#### Females

The female social defeat protocol described by Harris et al. (Harris et al., 2018) was used, with some adjustments. Screening of CD-1 mice and defeat sessions were carried out in the same way as for male, with the only difference being that female mice were covered with previously collected urine from a separate cohort of CD-1 male mice before each defeat session (**Fig. 1G**). This ensured that CD-1 mice displayed the required level of aggression towards females. The urine was applied using a paintbrush to the base of the tail, vaginal orifice, back and face of the female mice, using approximately 40µl per mouse (van Doeselaar et al., 2021). Each female received the same urine mixture, but the mixture varied daily to prevent the aggressors from becoming familiar with the scent. Immediately after urine application, the female was placed with the CD-1 aggressor. Defeat sessions lasted for either 5 minutes or 10 attacks but were ended immediately if mounting occurred. Aggressors that underperformed on more than one day were replaced with a new aggressor. Control females were housed in pairs and moved between cages every day. They had no physical contact with each other or CD-1 aggressor, did not have urine applied to them, and were housed in a different room to the defeated and CD-1 mice.

### Urine collection

Urine was collected from male CD-1 mice into 1.5ml Eppendorf tubes while manually restraining the mouse. Fresh urine was collected up to 1 week prior to use, mixed together and stored at +4°C.

### Body weight measurement

Body weights were measured before the CSDS exposure and daily throughout the social defeat period using standard laboratory scale.

### Behavioral tests

All behavioral testing was done during the light cycle. The tests were recorded and mice tracked using the automated video-tracking system ANY-maze (Stoelting Co., Wood Dale, Il). In case manual scoring was needed, it was performed by an experienced observer, blinded to the experimental condition.

#### Social Interaction (SI) Test

24 hours after the last defeat, the defeated mice were tested in the social interaction (SI) test. This was carried out in an open field (OF) box (45 × 45 × 45 cm) with a Plexiglas wire mesh enclosure (10 cm wide × 6.5 cm deep × 42 cm high) placed in a section of the box designated as the ‘social interaction zone’ (SIZ). In the first phase of this test, the mouse was placed in the box for 150 seconds with an empty wire mesh enclosure. The mouse was then placed back in the home cage for 30 seconds. For the second phase, the mouse was placed in the box for another 150 seconds, but this time with a novel CD-1 aggressor present in a different wire mesh enclosure. The time the mouse spent in the SIZ during each phase was measured and was used to calculate the individual’s SI ratio.

SI ratios were calculated by dividing the time spent in the SIZ with an aggressive CD-1 mouse present by the sum of time spent in the SIZ with and without the aggressor present (Henriques-Alves and Queiroz, 2016). This is a slightly adjusted version of the original equation (Golden et al., 2011) that allows the inclusion of mice that spent 0 seconds in the SIZ without the aggressor present. Mice subjected to social defeat were categorized into resilient (CSDS^RES^) and susceptible (CSDS^SUSC^) cohorts depending on their SI ratio, with a ratio of <0.5 being an index of susceptibility and a ratio of >0.5 being an index of resilience.

The time in the interaction zone with the target present and the SI ratio were used to calculate z-score in the SI test (zSI).

A second SI (SI(2)) test was carried out 24 hours after the last day of the LH protocol, to determine whether extinction of the CSDS resilient phenotype occurred.

#### Open Field (OF) Test

Locomotor activity was measured with a VersaMax animal activity monitor for 10 minutes in OF chambers (40 × 40 × 30 cm; Omnitech Electronics, Columbus, OH, USA) with photocells and plexiglass walls and floors. Total distance traveled (cm), time spent in the center and margin area of the OF, and vertical activity were recorded by the VersaMax system. The time in the center of the OF and ratio of distance traveled in the center (distance traveled in the center divided by the total distance traveled × 100) were used to calculate z-score in the OF (zOF). The total distance traveled was used to normalize for the locomotor component.

#### Elevated Plus Maze (EPM) test

The elevated plus maze (EPM) consisted of two opposing open arms (30 × 5cm) and two opposing closed arms (30 × 5 × 11cm) branching out from a center zone (5 cm^2^) elevated 50 cm above the floor. The mice were placed in the center zone of the EPM, facing an open arm, and were allowed to explore the maze for 10 minutes. The mouse was considered to be inside an open or closed arm when all four paws were inside it. The amount of time spent in the open arms and ratio of entries into the open arms (entries into open arms divided by total entries × 100) were used to calculate z-score in the EPM (zEPM). The total number of arm entries was used to normalize for the locomotor component.

#### Sucrose Preference Test

The sucrose preference test (SPT) was carried out as previously described (Liu et al., 2018), but in the animal’s home cage. Briefly, during the first 48 hours of habituation the mice were presented with two bottles along with regular food, one with water and second one with 1% sucrose solution, swapped each day to avoid place preference. On day 3, the bottles were removed from the cage, filled with fresh water and sucrose solution, and weighed for the baseline measure. The food was removed, and the water and sucrose bottles were placed in the cage. On day 4, the mice were given normal food and water until 8pm, the bottles were removed and weighed. The baseline sucrose preference measurement was calculated as a percentage of the weight of the sucrose bottle over the total weight of the sucrose and water bottles. All the steps were then repeated for the second baseline measurement, which took place on day 5. Mice had access to regular food and water until 8pm on day 5, and then underwent food and water deprivation for 24 hours. At 8pm on day 6, the mice were given fresh water and sucrose solution, after these bottles had been weighed. At 8am on day 7, the bottles were removed and weighed again. Sucrose preference was calculated as a percentage of the weight of the sucrose bottle over the total weight of the sucrose and water bottles.

#### Learned Helplessness (LH) test

The LH test was performed as previously described (Anisman and Merali, 2001). Briefly, in a two-compartments shuttle box (Imetronic, France) with an electrical grid floor, separated by an automatic gate, mice were first exposed to two training sessions 24 hours apart. Mice received pseudorandom electric footshocks (120 shocks, 0.25mA) over 30 minutes in one of the compartments, while the gate was closed. On the test day, 24 hours later, mice were subjected to an assessment session, which consisted of 20 shocks. Before the initial shock, there was a 3-minute habituation period during which the mouse could explore the two compartments. The gate between the compartments opened each time a shock occurred, giving the mouse the choice to escape to the other compartment. Each shock was terminated either as soon as the animal shuttled to the other compartment (defined as an escape), or at the end of the 24 second shock period (defined as a failure). Latency to escape each shock and the number of escapes and failures were recorded. Following the completion of test day 1, they were also subjected to assessment sessions on test days 10, 20 and 30, to investigate changes in helpless behavior over time. K-means (k=2) clustering analysis using the escape latency and number of escape failures for each individual was applied to categorize the mice into non-helpless (resilient, LH^RES^) or helpless (susceptible, LH^SUSC^) for each test day.

### Cumulative Stress-sensitivity Indexes Calculation

Z-score analysis was used to give each mouse its own individual score of stress sensitivity (similar to (Guilloux et al., 2011)) by integrating the data from each behavioral test (SI test, OF test, EPM test), along with change in weight during CSDS. Briefly, z-score values were calculated for test parameters by subtracting the mean value of the sex-matched control from each test individual’s value and then divided by the control group’s standard deviation. The directionality of scores was adjusted so that increased score values reflected increased dimensionality. The resulting indexes were averaged across tests to generate an individual cumulative stress-sensitivity index, which was then averaged within each group.

### Brain Tissues Collection

24 h after the last behavioral test mice were anaesthetized individually using isoflurane before being sacrificed by rapid decapitation. The brain tissue was then extracted and placed immediately in an isopentane solution resting in dry ice, at a temperature between -30 and -40 C. The frozen brain tissues were then stored in a -80 freezer until needed.

On the day of dissection, the brain was placed in a cryostat (Leica CM3000, Leica Biosystems, France) for coronal slicing. Using a magnifying glass, the discrete regions (NAc, AMY, VTA) were identified with the reference of a mouse brain atlas (Franklin and Paxinos, 2008), taken out from the brain slices (left and right hemispheres) using punch pens (Harris Uni-Core) of different inner diameters (1 mm, 0.75 mm, 0.5 mm) and collected in labelled 1.5 mL Eppendorf tubes. The tissue samples were stored at −80 °C until further western blot analysis.

### Western Blot

The tissue punches from 3 mice were pooled per sample and homogenized in RIPA lysis buffer on ice. The homogenates were centrifuged at 10,000×*g* for 30 min and the supernatant used for western blot analyses. Pierce™ BCA Protein Assay kit (Thermo Scientific, Rockford, IL, USA) was used for protein concentration measurement, and 20 µg of protein in Laemmli sample buffer with a final concentration of 10% β-mercaptoethanol were boiled in a water bath for 5 min. The proteins were separated by 12% SDS-polyacrylamide gel electrophoresis and transferred to nitrocellulose membrane. The membrane was blocked with 5% BSA (A9647, Sigma-Aldrich, MO, USA) for 1 h and then incubated overnight at 4°C with primary antibodies: GAPDH (1:500; sc-32233), cFos (1:250; sc-52-G), pERK (1:500; sc-7383) from Santa Cruz Biotechnology, Inc., ERK (1:500; 9102S, Cell Signaling Technology, Inc.), and BDNF (1:500; AB108319, Abcam). After 24 h, membranes were rinsed three times with TBST and incubated with secondary antibodies IRDye IR800 goat anti-rabbit (1:2500; 611-132-122) and IRDye IR700 goat anti-mouse IgG (1:2500; 610-130-121) from Rockland Immunochemicals (Limerick, PA, USA) for 2 h at RT. Membranes were visualized and protein bands were quantitatively analyzed using the LI-COR Odyssey Imaging System (LI-COR Biotech LLC). The results are given as ratio of the optical density of the protein of interest to the corresponding GAPDH band.

### Statistical Analyses

Statistical analysis was performed using the Prism software (version 10, GraphPad Software, San Diego, CA, USA). Normality of distribution of the data was assessed using the Shapiro-Wilk test. When at least one group in a given comparison failed the normality test, nonparametric alternatives were used. For the comparisons of two independent groups unpaired Student’s t-test was used. Behavioral parameters with multiple group comparisons were analyzed either with one-way ANOVA (normal distribution), followed by Tukey’s multiple comparisons test to compare individual groups when a main effect was detected or Kruskal-Wallis (non-normal distribution), followed by Dunn’s multiple comparisons test. The Fisher’s exact test was used to compare the proportions of observations between groups. All data are presented as mean ±SD. Differences were considered significant at *p* < 0.05.

## Results

### 1. Chronic social defeat stress induced avoidance behavior in both males and females

To investigate cross-resilience to social defeat stress and learned helplessness we carried out several behavioral tests displayed in the experimental timeline (**Fig. 1A**), beginning with the CSDS model, since CSDS phenotype lasts up to one month, unlike LH (Meduri et al., 2013). In the defeated group, male (n=38) and female (n=23) mice were subjected to 10 days of CSDS, which was followed by a social interaction (SI) test for both defeated animals and controls (males n=30, females n=18) to screen for social avoidance behavior (**Fig. 1B, G**).

In Males, 42% of the defeated males were categorized as resilient (CSDS^RES^) and 58% as susceptible (CSDS^SUSC^) (**Fig.1C**), which validates the social defeat protocol as 30-40% of males are expected to show a resilient phenotype (Golden et al., 2011). CSDS^SUSC^ males exhibited increased social avoidance, spending significantly less time in the SIZ when the CD-1 mouse was present, compared to the controls (CSDS^CTR^)(**Fig.1D**: Kruskal-Wallis test, H(2) = 15.41, P=0.0005; **Suppl. Fig.1A**). Dunn’s multiple comparisons test showed significant difference between CSDS^CTR^ vs CSDS^SUSC^ (adj.p=0.0003), and trend to significance between CSDS^SUSC^ vs CSDS^RES^ (adj. p=0.0663). Kruskal-Wallis test showed that there was a significant difference in the SI ratios between the CSDS^CTR^, CSDS^SUSC^, and CSDS^RES^ cohorts (**Fig.1E**: H(2)=26.77, P<0.0001). Dunn’s multiple comparisons test revealed significant difference between CSDS^CTR^ vs CSDS^SUSC^ (adj.p<0.0001), and CSDS^SUSC^ vs CSDS^RES^ (adj. p<0.0001), with CSDS^SUSC^ males having the lowest SI ratio. The analysis of sucrose intake did not show significant difference among CSDS^CTR^, CSDS^SUSC^, and CSDS^RES^ males (**Suppl. Fig.2A**).

**Figure 2.**
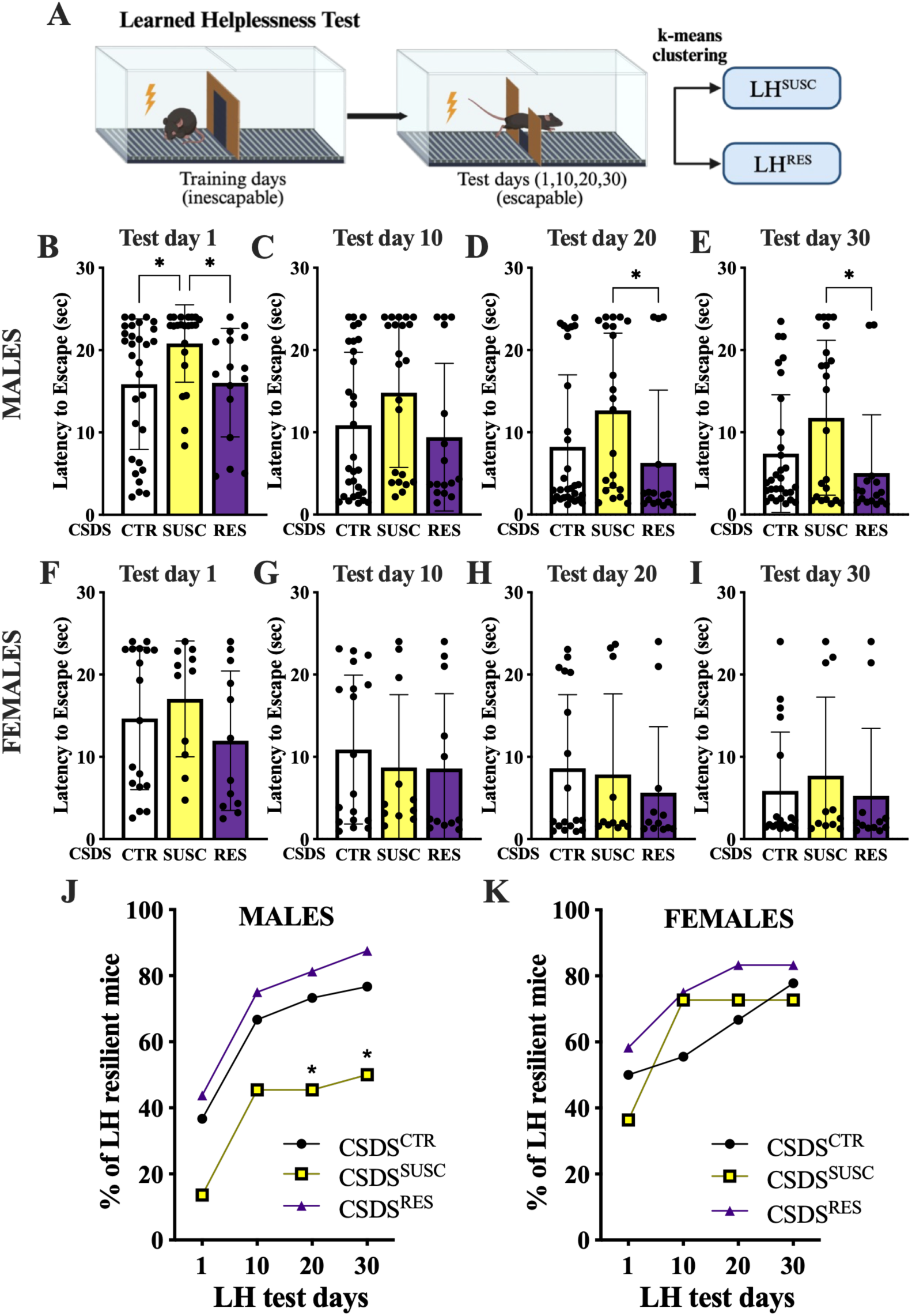
Effect of social defeat stress on escape behavior in learned helplessness test. A. Graphic representation of LH paradigm (the image was created in BioRender). The latency to escape for each LH test day for males (B-E) and for females (F-I). The proportion of mice that were LH^RES^ for CSDS^CTR^, CSDS^SUSC^ and CSDS^RES^ groups for males (J, \**p*<0.05 vs CSDS^RES^) and females (K). Orange lines on spider plots indicate the proportion of mice RES to LH test, and blue lines – proportion of SUSC mice. Data are shown as mean ± SD, with \**p*<0.05 indicating significance.

In females, 52.2% of the defeated female mice were categorized as resilient and 47.8% as susceptible (**Fig.1G**), percentages comparable to those obtained by other researchers using the same female social defeat protocol (Harris et al., 2018; Ortiz et al., 2022). Similar to males, CSDS^SUSC^ females exhibited increased social avoidance, spending significantly less time in the SIZ when the CD-1 aggressor was present compared to the control and resilient groups (**Fig.1H**: One way ANOVA, F(2,38) = 7.181, P=0.0023; **Suppl.Fig.1B**). Tukey’s multiple comparisons test showed significant difference between CSDS^CTR^ vs CSDS^SUSC^ (adj. p= 0.0062), and CSDS^SUSC^ vs CSDS^RES^ (adj. p=0.0041). Also, there was a significant difference between the SI ratios of the CSDS^CTR^, CSDS^SUSC^, and CSDS^RES^ cohorts, and susceptible females had the lowest SI ratios (**Fig.1I**: Kruskal-Wallis test, H(2) = 17.72, P=0.0001). Dunn’s multiple comparisons test revealed significant difference between CSDS^CTR^ vs CSDS^SUSC^ (adj.p=0.0003), and CSDS^SUSC^ vs CSDS^RES^ (adj.p=0.0013). Similar to males, females did not show differences among groups in sucrose intake (**Suppl. Fig.2B**).

To investigate differences in individual stress vulnerability between CSDS resilient, susceptible and control groups, we gave each mouse its own individual score of stress-sensitivity based on individual behavior and physiological parameters (**Suppl. Fig.3-5; Suppl. Table 1**), as described in the Methods. In males, one-way ANOVA test revealed significant difference between groups (**Fig.1F**: F(2,65) = 10.29, P=0.0001). *Post hoc* Tukey’s test indicated a significant difference in cumulative indexes between CSDS^CTR^ vs CSDS^SUSC^ (adj.p=0.0001), and CSDS^RES^ vs CSDS^SUSC^ (adj.p=0.0086) groups. As observed in males, females also exhibited significant difference in cumulative indexes between groups (**Fig.1K**: one-way ANOVA test, F(2,38) = 16.72, P<0.0001). Tukey’s multiple comparisons test showed significant difference between CSDS^CTR^ vs CSDS^SUSC^ (adj. p<0.0001), and CSDS^RES^ vs CSDS^SUSC^ (adj.p=0.0047). This data suggests that for both males and females CSDS^SUSC^ mice have higher overall vulnerability to stress.

**Figure 3.**
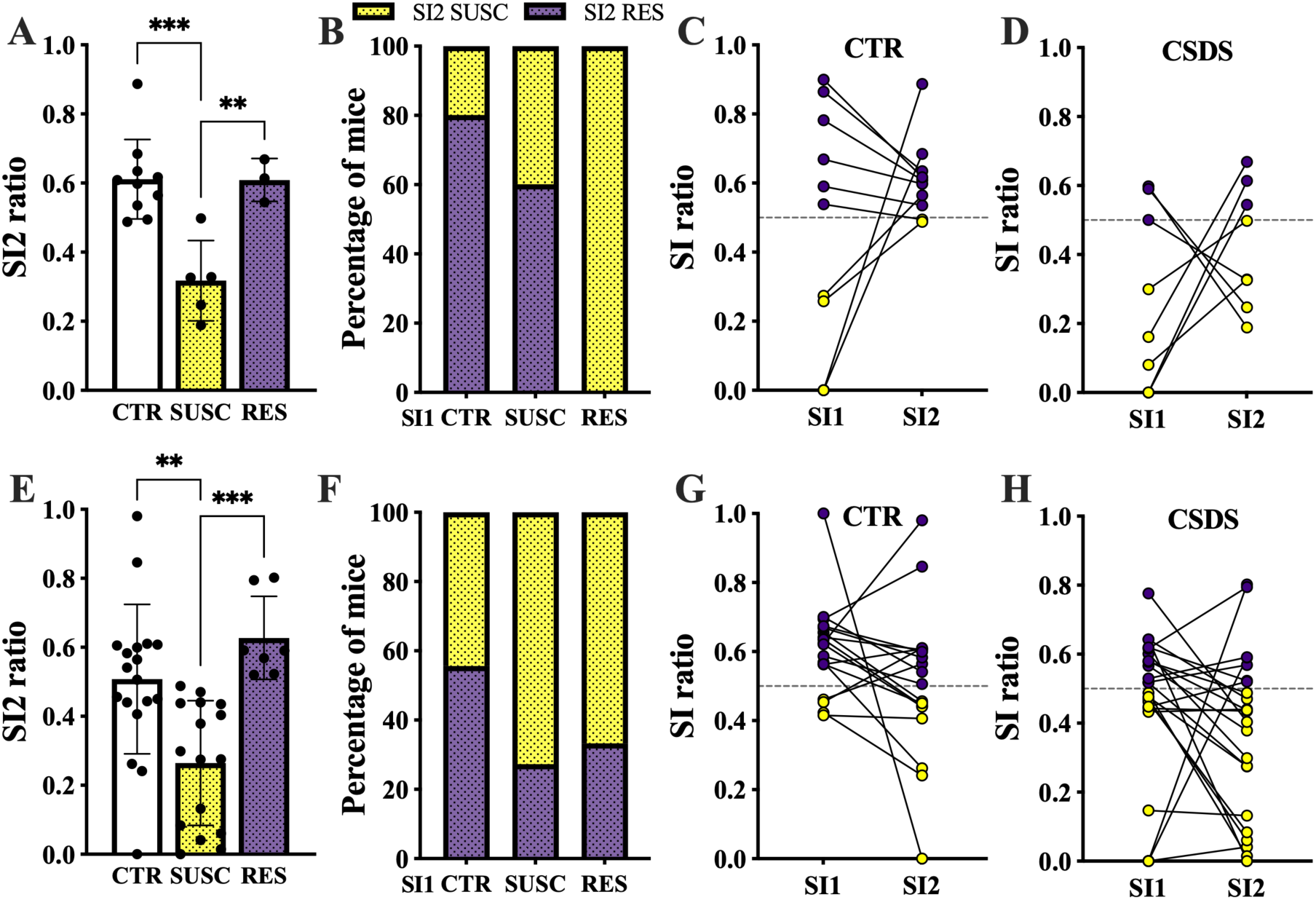
Effect of learned helplessness test on social interaction of previously categorized SUSC, RES and CTR male and female mice. Social interaction ratio in the second social interaction test, carried out after the last learned helplessness test day for males (A) and females (E). The percentage of mice being RES or SUSC in the second SI test for RES, SUSC and CTR males (B) and females (F) in the initial SI test. Changes in SI ratio between the initial SI and the second SI for CTR males (C) and females (G) and CSDS exposed males (D) and females (H). Data are shown as mean ± SD, with \**p*<0.05, \*\**p*<0.01, ***p<0.001 indicating significance.

### 2. Resilience to CSDS predicts resilience to LH in males, but not in females

Cross-resilience to social defeat stress and learned helplessness was assessed by subjecting defeated and control CSDS groups to a LH uncontrollable foot-shock stress protocol (**Fig 2A**). The foot-shock escape behavior (number of failures to escape and latency to escape) was assessed and mice were classified into stress-resilient (non-helpless) or stress-susceptible (helpless) using k-means clustering algorithm.

The average latency to escape foot-shocks on test day 1 was significantly lower in CSDS^RES^ males as compared to CSDS^SUSC^, and CSDS^SUSC^ mice had a significantly higher latency to escape than CSDS^CTR^ (**Fig. 2B**: Kruskal-Wallis test, H(2)=10.52; P=0.0052 for main group effect; CSDS^CTR^ vs CSDS^SUSC^: adj.p=0.0147, SUSC vs RES: adj.p=0.0158). There were no significant differences between groups for latency to escape on test day 10 (**Fig.2C**), however on test day 20 and test day 30 CSDS^RES^ mice had a significantly lower latency to escape the foot-shocks than CSDS^SUSC^ mice (TD20: Kruskal-Wallis test, H(2)=7.252, P=0.0266 for main group effect; CSDS^SUSC^ vs CSDS^RES^ adj.p=0.0242 (**Fig. 2D**); TD30: Kruskal-Wallis test, H(2)=5.939, P=0.0513 for main group effect; CSDS^SUSC^ vs CSDS^RES^ adj.p=0.0469 (Fig. 2E)). In females, for CSDS^RES^ it took less time to escape the foot-shocks than CSDS^SUSC^ females, but there wasn’t a significant difference between groups on none of the test days (**Fig.2F-I**).

To further investigate cross-resilience to chronic social stress and learned helplessness, we calculated the proportions of individuals that were resilient on each LH test day for CSDS^RES^, CSDS^SUSC^ and CSDS^CTR^ groups. In males, on the LH test day 1, the proportion of LH resilient (LH^RES^) mice among CSDS resilient (CSDS^RES^®LH^RES^) was 43.8% (**Fig.2J, Suppl.Fig.6A-C**), whereas among CSDS susceptible (CSDS^SUSC^®LH^RES^) it was only 13.6% (Fisher’s test, p=0.0623). By LH test day 10 the proportion of CSDS^RES^®LH^RES^ increased to 75%, compared to 45.6% in CSDS^SUSC^®LH^RES^ (Fisher’s test, p= 0.1). While the proportion of LH^RES^ mice did not increase much further in the CSDS^SUSC^ group, it increased in CSDS^RES^ group, reaching 81.3% on test day 20 (Fisher’s test, p<0.05) and 87.5% on test day 30 (Fisher’s test, p<0.05). No significant differences were found in the proportion of LH^RES^ mice between the CSDS^RES^ and CSDS^CTR^ groups, suggesting that: 1) resilience to one stressor can protect against the adverse behavioral effect of another stressor similar to control individual who did not experience the initial stressor, 2) susceptibility to CSDS increases the likelihood of subsequent susceptibility to other stressors, 3) CSDS^SUSC^ mice demonstrate lower extinction of LH-helpless phenotype.

There appears to be a different relationship between resilience to social defeat and LH stress in females. Although there was a trend toward a higher proportion of LH^RES^ among CSDS^RES^ compared to CSDS^SUSC^ on test day 1, the proportions of LH^RES^ mice did not differ significantly among the CSDS^CTR^, CSDS^SUSC^ and CSDS^RES^ groups on test day 1, 10, 20 and 30 (**Fig. 2K, Suppl.Fig.6D-F**). This suggests that, unlike in males, CSDS susceptibility in females is not strongly associated with the likelihood of subsequent susceptibility to foot-shock stress.

In addition, comparison between males and females revealed that 36.4% of females CSDS^SUSC^ were LH^RES^ on LH test day 1, compared to only 13.6% in males (p=0.1863). Moreover, by the final LH test day, 72.7% of the CSDS^SUSC^ females were LH^RES^, compared to 50% of the CSDS^SUSC^ males, (p=0.2783). While the differences were not significantly different, it may suggest that females vulnerable to chronic social stress are less likely to also be vulnerable to trauma-type stress compared to males.

Resilient and susceptible phenotypes in the CSDS model are known to persist for at least 30 days after the defeat (Meduri et al., 2013). To determine whether engagement in the learned helplessness (LH) test alters the maintenance or extinction dynamics of CSDS-induced susceptibility, we performed a second SI test 24 hours after the day-30 LH session. Following the second interaction test (SI(2)), mice were reassigned into new CSDS^SUSC^ and CSDS^RES^ groups based on their SI(2) ratios.

In males, the second SI test revealed clear group differences in social interaction behavior (**Fig. 3A**; one-way ANOVA, F(2,15) =12.89, P=0.0006). Mice newly classified as CSDS^SUSC^ showed significantly lower SI2 ratios compared with both CSDS^CTR^ (adj.p=0.0005) and CSDS^RES^ animals (adj.p=0.0065). Surprisingly, 100% of mice that were CSDS^RES^ following the first SI test were categorized as CSDS^SUSC^ in the second SI test, and 60% of the mice that were CSDS^SUSC^ following the first SI test were categorized as CSDS^RES^ in the second SI test (**Fig. 3B-D**). This data suggests that being exposed to trauma-type stress such as foot-shocks changes an individual’s response and vulnerability to social defeat stress in males.

In females there was also a significant difference between the SI2 ratios of the CSDS^CTR^, CSDS^SUSC^, and CSDS^RES^ groups (**Fig.3E**; One-way ANOVA, F(2, 38)=11.29, P=0.0001). Females newly categorized as CSDS^SUSC^ in the second SI test had significantly lower SI2 ratios compared with both CSDS^CTR^ (adj.p=0.0018) and CSDS^RES^ (adj.p=0.0004). 66.7% of females that were CSDS^RES^ following the first SI test were categorized as CSDS^SUSC^ in the second SI test, and 27.3% of the mice that were CSDS^SUSC^ following the first SI test were categorized as CSDS^RES^ in the second SI test (**Fig. 3F-H**). This finding differs from males, indicating sex differences in how LH exposure affects the retention of prior CSDS experience.

### 3. Mice that are resilient or susceptible in both tests demonstrate sex- and region-specific differences in ERK pathway protein expression

We assessed the levels of BDNF, total ERK1/2, and phospho-ERK1/2 in the ventral tegmental area (VTA), amygdala (AMY), and nucleus accumbens (NAc) of CSDS^RES^®LH^RES^ (RES) and CSDS^SUSC^®LH^SUSC^ (SUSC) mice. In males, BDNF levels were significantly lower in the SUSC group compared to the RES group in both the NAc (t-test, t=3.788, p<0.05, Fig. 4A) and AMY (t-test, t=5.724, p<0.01, Fig. 4C). In contrast, females exhibited significantly higher BDNF levels in the SUSC group than in the RES group in the VTA (t-test, t=5.886, p<0.01, Fig. 4F). Phospho-ERK1/2 levels were significantly lower in the VTA of SUSC males compared to their RES counterparts (t-test, t=5.112, p<0.05, Fig. 4E). In females, total ERK1/2 levels were significantly decrease in the AMY (t-test, t=5.659, p<0.05, Fig. 4D), and increase in the VTA of SUSC mice (t-test, t=4.225, p<0.05, Fig. 4F) compared to RES females. These findings highlight sex-specific differences in BDNF-ERK signaling pathway within key brain regions associated with response to stress, reward and emotional experiences.

**Figure 4.**
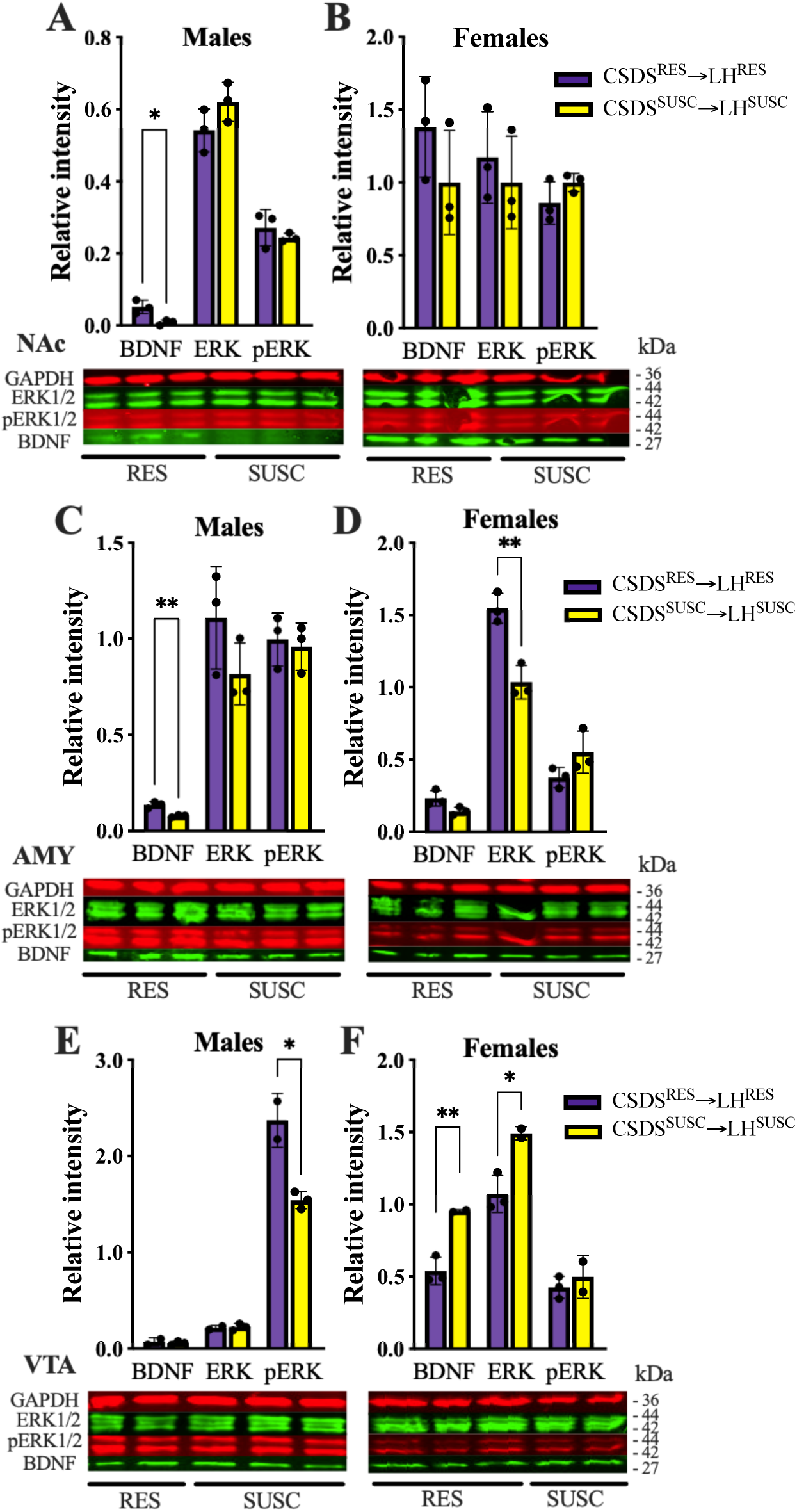
Western blot of phospho-ERK, total ERK and BDNF in CSDS RES and SUSC mice. Quantitative data and representative western blots demonstrating the levels of analyzed proteins in NAc of males (A) and females (B), in AMY of males (C) and females (D), and in VTA of males (E) and females (F). Each data point represents the pooled brain tissues extracted from three mice. Data are shown as mean ± SD, with \**p*<0.05, \*\**p*<0.01 indicating significance.

### 4. NEKO females more effectively retain the CSDS resilient phenotype following learned helplessness exposure compared to WT mice

In our preceding study (Isingrini et al., 2016), we demonstrated that NE neurotransmission regulate vulnerability to chronic social defeat stress. Using conditional *VMAT2^DBHcre^* knockout (NEKO) mice (**Fig 5A**), we previously showed that only a very small proportion of NEKO defeated male mice displayed a resilient phenotype (5.3%). Because NEKO males are not suitable for cross-resilience studies due to their lack of resilience to CSDS, we only used NEKO females in the subsequent cross-study.

**Figure 5.**
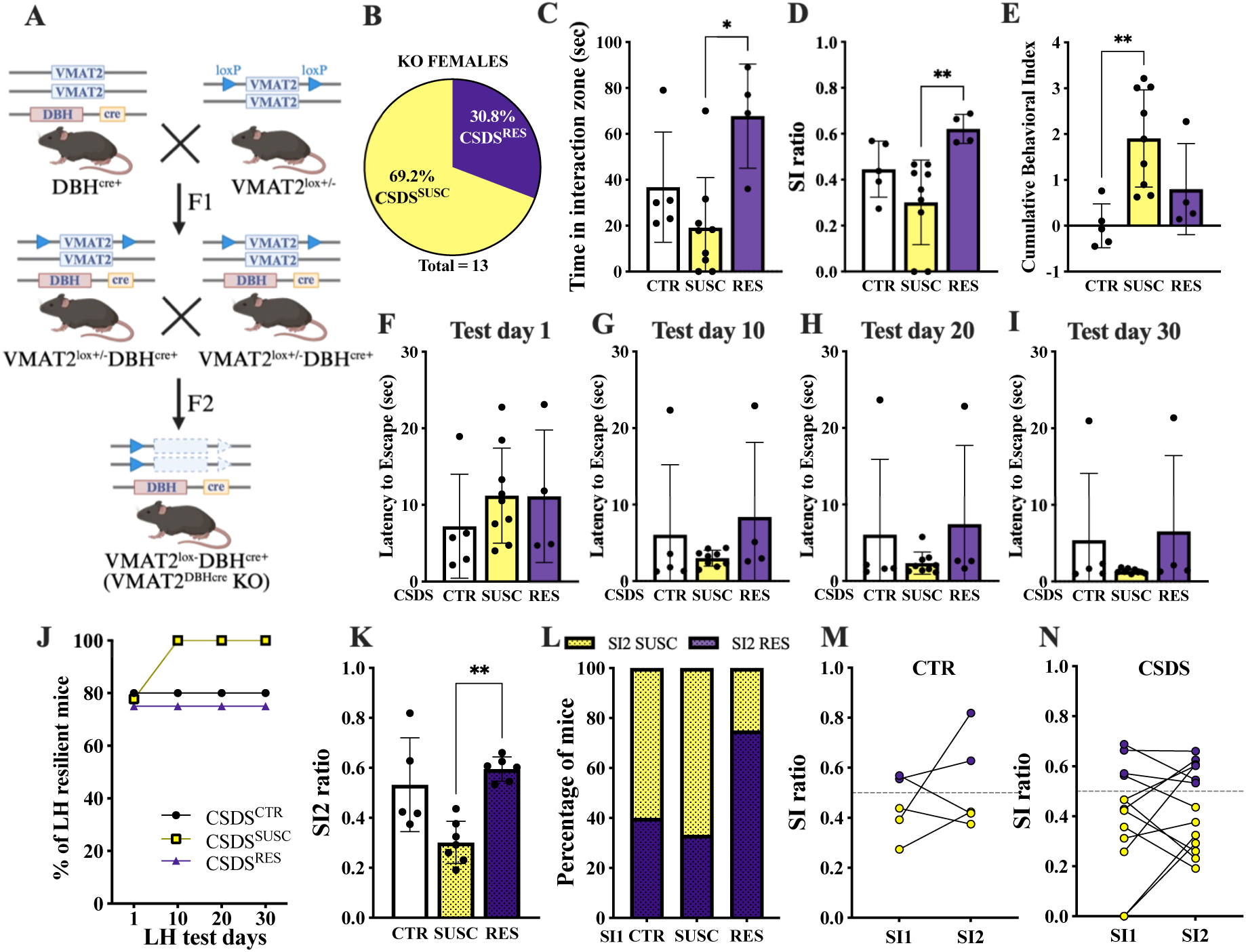
Behavior of NE-KO females stressed and control groups. A. Schematic diagram of generating a conditional knockout (cKO) mouse from breeding a DBH-cre mouse (top left) with a mouse carrying heterozygous floxed VMAT2 gene (top right). The image was created in BioRender. B. The proportion of CSDS resilient and susceptible KO females. C. The time spent in the interaction zone when the target mouse was present. D. Social interaction ratio. E. Susceptibility index. F-I. The latency to escape for each LH test day. J. The proportion of mice that were LH^RES^ for CSDS^CTR^, CSDS^SUSC^ and CSDS^RES^ groups. K. Social interaction ratio in the second social interaction test carried out after the last learned helplessness test day. L. The percentage of mice being RES or SUSC in the second SI test for RES, SUSC and CTR. Changes in SI ratio between the initial SI and the second SI for CTR (M) and CSDS exposed (N). Data are shown as mean ± SD, with \**p*<0.05, \*\**p*<0.01 indicating significance.

NEKO female mice yielded 30.8% of resilient individuals when tested in the SI test following CSDS paradigm **(Fig. 5B)**, therefore more likely to be susceptible to CSDS than WT females, that showed 52.2% RES. Indeed, CSDS^SUSC^ NEKO females demonstrated increased social avoidance, spending significantly less time in the SIZ with the CD-1 aggressor compared to the CSDS^RES^ NEKO (**Fig 5C**: Kruskal-Wallis test, H(2) = 8.511, P=0.0067; **Suppl. Fig.1C**). Dunn’s multiple comparisons test revealed significant difference between CSDS^SUSC^ vs CSDS^RES^ (adj.p=0.0138). Kruskal-Wallis test identified a significant difference between the SI ratios among the CSDS^CTR^, CSDS^SUSC^, and CSDS^RES^ cohorts (**Fig. 5D**: H(2) = 9.587, P=0.0025). Subsequent Dunn’s multiple comparisons test confirmed significant difference in SI ratios between CSDS^SUSC^ vs CSDS^RES^ (adj.p=0.0059). When comparing the cumulative behavioral indexes, Kruskal-Wallis test revealed significant differences among the groups (H(2) = 9.998, P=0.0016), with Dunn’s *post hoc* showing that CSDS^SUSC^ NEKO females had higher cumulative behavior index compared to CSDS^CTR^ NEKO females (**Fig. 5E**: adj.p=0.0016).

In learned helplessness test, the average latency to escape for NEKO females did not differ between groups on any LH test days (**Fig. 5F-I**). The proportion of CSDS^RES^®LH^RES^ and CSDS^CTR^®LH^RES^ did not change throughout the LH test and amounted to 75% and 80% respectively, however, CSDS^SUSC^®LH^RES^ increased from 77.8% on test day 1 to 100% on all subsequent test days (**Fig. 5J; Suppl.Fig.6G-I**). The proportion of LH^RES^ mice did not differ significantly between the groups.

NEKO females subjected to the second SI test 24 hours after LH test day 30 demonstrated significant difference in SI2 ratios between CSDS^CTR^, CSDS^SUSC^, and CSDS^RES^ groups (**Fig. 5K**: Kruskal-Wallis test, H(2) = 10.25, P=0.0017), with newly classified CSDS^SUSC^ NEKO females having significantly lower SI2 ratios than CSDS^RES^ KO (Dunn’s test, adj.p = 0.0066). Only 33.3% of the mice initially categorized as CSDS^SUSC^ after the first SI test were reclassified as CSDS^RES^ during the second SI test (**Fig. 5L**), like what was observed in WT females. Nonetheless, unlike WT females, 75% of KO female mice classified as CSDS^RES^ after the first SI test remained CSDS^RES^ following the second SI test. This suggests that in NE-depleted mice, foot shock did not markedly reverse the CSDS^RES^ phenotype but instead contributed to its preservation during the second SI test.

## Discussion

Depressive-like behavior assessed in rodents is typically evaluated through a series of behavioral tests covering several aspects of symptoms, analogous to those diagnosed in humans. These assessments may involve analyzing individual behavioral tests separately or conducting a comprehensive analysis that integrates multiple behavioral parameters from various tests. In our study, to evaluate depression-related phenotypes following CSDS exposure, we employed commonly used behavioral tests, including social avoidance and anxiety-like behavior, alongside physiological measures such as body weight fluctuations. While some studies have reported an increased tendency for anxiety-like behavior following social defeat stress (Kinsey et al., 2007; Krishnan et al., 2007; Yin et al., 2019; Matsumoto et al., 2021), our findings align with studies that did not observe significant differences in anxiety-like behavior (Jianhua et al., 2017; Liu et al., 2017) or anhedonia (Henriques-Alves and Queiroz, 2015; Matsumoto et al., 2021). However, we identified significant alterations in avoidance behavior. To achieve a more comprehensive evaluation, we applied a generalized approach using z-normalization, which confirmed that both male and female mice exhibiting susceptible (CSDS^SUSC^) phenotype had a higher cumulative index than control (CSDS^CTR^) or resilient (CSDS^RES^) animals. Together, our findings suggest that CSDS induces common endophenotypes of major depressive disorder, particularly social avoidance, and that CSDS^SUSC^ animals were significantly more sensitive to stress overall.

MDD or PTSD are highly comorbid, with approximately half of PTSD patients also suffering from MDD (Kessler et al., 1995). Conversely, individuals with MDD are at higher risk of developing PTSD after exposure to a traumatic event (Koenen et al., 2002). However, there is limited research that considers resilience to both chronic and trauma-type stress within the same individual, particularly in preclinical models. As a result, very little is known about the association of resilience to both types of stress. In this context, our current results provide new insight, indicating that there is cross-resilience to both CSDS and LH stress in male mice. We found that a higher proportion of males that were resilient following 10 days of social defeat were also resilient when tested 30 days following LH foot-shock stress, compared to males that were susceptible following social defeat. By the final test days, this difference in proportion was significant, indicating that CSDS^RES^ individuals exhibit faster extinction of the LH susceptible phenotype. These results suggest that males that are resilient to chronic social stress are more likely to be resilient to trauma-type stress, which supports our hypothesis that there is a cross-resilience. In addition, we found that CSDS^SUSC^ males had a higher cumulative behavior index than CSDS^RES^ males and controls. The difference in indexes suggests that mice that are susceptible to chronic social stress have greater overall stress vulnerability and those that are resilient have the lowest, which provides further support for the cross-resilience hypothesis.

Some studies suggest the existence of similar mechanisms underlying stress-susceptibility in males and females. For instance, Ortiz et al. (Ortiz et al., 2022) showed that susceptibility to stress in females can be induced through the positive modulation of α7 nicotinic acetylcholine receptors (nAChRs), an effect previously observed in male mice (Morel et al., 2018). However, other research highlights sex differences in the mechanisms of resilience. As an example, (Curtis et al., 2006) found that in females, lower levels of corticotropin-releasing factor (CRF) are required to activate the locus coeruleus-norepinephrine (LC-NE) arousal system following stress exposure. Our results demonstrate that there are sex differences in the patterns of resilience to chronic social and trauma-type stress. While there were a higher proportion of CSDS^RES^ females that were resilient to LH on each test day compared to CSDS^SUSC^ females, there were no significant differences in latencies to escape. This suggests that vulnerability to chronic social stress does not predict vulnerability to trauma-type stress in females, so they may not have the same pattern of cross-resilience as males. A higher proportion of CSDS^SUSC^ females were resilient to LH on each test day compared to CSDS^SUSC^ males, which could indicate that females are more likely to be resilient to trauma-type stress than males. In addition, females did not show the same phenotype reversal in the second SI test that the males did, suggesting there are sex differences in extinction of CSDS susceptibility or resilience. On test days 10 and 20 of LH, a higher proportion of CSDS susceptible females were resilient compared to controls, which is something we did not observe in males and could indicate that females that undergo chronic social stress have greater resilience to trauma-type stress than those that have not previously been exposed to stress. However, in contrary to this, the cumulative indexes of CSDS susceptible females were significantly higher than those of the controls, which suggests that CSDS susceptible females are more vulnerable to stress overall.

The distinct molecular adaptations observed in males and females suggest that different signaling cascades contribute to resilience, likely reflecting sex-specific neurobiological strategies for coping with stress exposure. In males, cross-resilience correlated with elevated BDNF and phospho-ERK levels in the Nacc and VTA, two regions integral to motivation and reward processing. This aligns with prior evidence implicating mesolimbic plasticity in resilience and suggests that dopaminergic regulation is a key component of stress adaptation in males. This is consistent with previous studies that reported stress-induced increases in phospho-ERK in VTA (Iñiguez et al., 2010; Yap et al., 2015), suggesting that elevated phospho-ERK levels in resilient males may enhance motivation, thereby facilitating stress adaptation. BDNF primarily signals through the TrkB receptor, activating intracellular cascades such as the ERK/MAPK, PI3K/Akt, and PLCγ pathways (Numakawa and Kajihara, 2023). The lack of total ERK changes in males suggests that BDNF-mediated resilience may involve alternative signaling branches, such as PI3K/Akt, which promotes neuronal survival and synaptic plasticity, or PLCγ, which regulates calcium-dependent processes critical for neuroplasticity.

Conversely, females showed region-and phenotype-specific changes in BDNF and ERK signaling, but these did not align with a clear behavioral cross-resilience profile. In susceptible females, increased BDNF in the VTA coupled with decreased ERK in the amygdala suggests a distinct neuroadaptive strategy potentially oriented around emotion and arousal regulation. The significant ERK and BDNF decrease in the VTA of resilient females may also suggest reduced TrkB-ERK signaling in the VTA that would limit activation of the mesolimbic dopamine system. Furthermore, sex hormones may interact with these molecular pathways to shape resilience. Estrogen has been shown to modulate ERK signaling and synaptic plasticity in a region-dependent manner (Bulayeva et al., 2004; Sarkar et al., 2010), potentially contributing to the sex-specific ERK regulation observed in resilient females. Additionally, glucocorticoid signaling, which is critical in stress responses, may differentially impact ERK and BDNF pathways across sexes. In males, glucocorticoid receptors (GRs) in the NAc and AMY may interact with BDNF-TrkB signaling to facilitate resilience, whereas in females, GR activation may influence ERK dynamics in the AMY and VTA.

Noradrenergic projections to the VTA and AMY are known to originate primarily from the locus coeruleus (LC), where they modulate neuron excitability and stress responses (Guiard et al., 2008; McCall et al., 2017). Notably, our previous work demonstrated that activation of NE signaling to the VTA enhanced resilience to CSDS in WT males, suggesting a critical role for NE in facilitating adaptive plasticity in this region (Isingrini et al., 2016). However, NE innervation of target brain regions may differ between males and females, with evidence showing that females have fewer noradrenergic cells but a larger dendritic volume than males (Mariscal et al., 2023). Considering that NE plays a role in regulating ERK/MAPK signaling (Tolbert et al., 2003; Price et al., 2004), the absence of NE signaling in *VMAT2^DBHcre^* knockout females could lead to failure to engage these molecular cascades, preventing stress-induced plasticity and behavioral adaptations that would typically shift individuals from resilience to susceptibility, and maintaining resilience despite additional stress exposure. However, whereas in males NEKO, there was almost no resiliency, about 30% of females NEKO are resilient, underlying a different role of NE transmission in males versus females. Together, these findings underline the essential role of NE in orchestrating stress-related neural adaptations and may trigger distinct resilience mechanisms in males and females, with females that may be supported by a more constrained or finely tuned neurochemical environment.

Future research should explore how sex hormones, developmental stress exposure, and context-specific factors shape resilience trajectories. Multi-omic approaches combined with neural circuit interrogation may help identify biomarkers of cross-resilience and clarify whether it represents a stable trait or a modifiable state.

In summary, our work introduces cross-resilience as a novel framework to study individual variability in stress response. We show that resilience to chronic social defeat can buffer against subsequent helplessness in males but not females, and that these differences are mirrored by distinct molecular signatures. Understanding the sex-specific architecture of resilience may lead to better-targeted interventions for stress-related disorders.

## Acknowledgements

The authors express their gratitude for the financial support of the CIHR to BG (Grant # 201909PJT-419517), Ferring Postdoctoral Fellowship in Health to MB, FRQS Postdoctoral Fellowship to GF. Mice breeding and genotyping was conducted by Ms Yeqing Geng.

## Notes

### Competing Interest Statement

The authors have declared no competing interest.

