## Supplementary Figures 1-6 for "Sex-Dependent Cross-Resilience to Social Defeat and Learned Helplessness: The Role of BDNF-ERK Signaling and Norepinephrine"

**Supplementary Figure 1** - Social interaction test: time in the interaction zone (IZ).

**Supplementary Figure 2** - Sucrose preference test.

**Supplementary Figure 3** - Open Field test (OF).

**Supplementary Figure 4** - Elevated Plus Maze test (EPM).

**Supplementary Figure 5** - Weight change after CSDS

**Supplementary Figure 6** - Radar plots representing the proportions of $LH^{RES}$ and $LH^{SUSC}$ animals

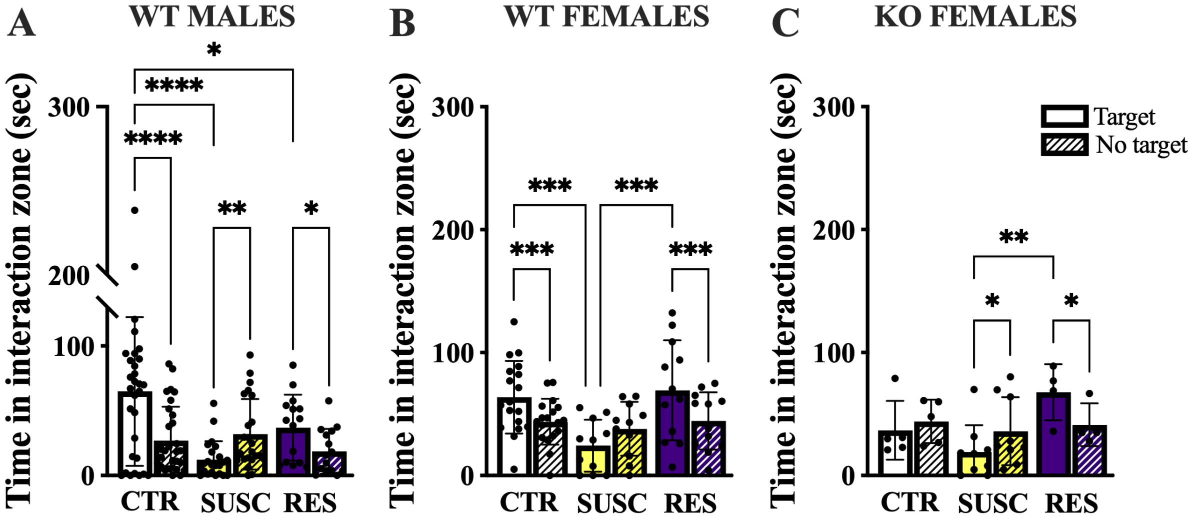


**Supplementary Figure 1 -** Social interaction test: time in the interaction zone (IZ).

In WT males (A), a two-way ANOVA revealed significant main effect of target (F(1,65)=9.828, P=0.0026), behavioral phenotype (F(2,65)=4.463, P=0.0153), and target x phenotype interaction (F(2,65)=22.75, P<0.0001). Tukey's *post hoc* analysis showed that $CSDS^{CTR}$ (adj.p<0.0001) and $CSDS^{RES}$ (adj.p=0.0248) males spent significantly more time in the IZ in the presence of CD1 mice (target) compared to the no-target phase. In contrast, time spent in the IZ with the target was significantly decreased in $CSDS^{SUSC}$ males (adj.p=0.0034). In WT females (B), a two-way ANOVA likewise revealed significant main effects of target (F(1,38)=8.125, P=0.0070), behavioral phenotype (F(2,38)=3.968, P=0.0272), and a target × phenotype interaction (F(2,38)=10.06, P=0.0003). Tukey's *post hoc* test showed that, similar to males, $CSDS^{CTR}$ (adj.p=0.0006) and $CSDS^{RES}$ (adj.p=0.0005) females spent significantly more time in the IZ during target compared to no-target sessions. $CSDS^{SUSC}$ females showed a trend toward spending less time in the IZ during target compared to no-target sessions (adj.p=0.0523), and they spent significantly less time in the IZ with the target compared to $CSDS^{RES}$ females (adj.p=0.0004). In KO females (C), a two-way ANOVA showed significant target × phenotype interaction (F(2,15)=6.606, P=0.0088). Tukey's *post hoc* test showed that $CSDS^{RES}$ KO females spent significantly more time in the IZ in the presence of target compared to the no-target phase (adj.p=0.0180), while $CSDS^{SUSC}$ KO females spent significantly less time in the IZ with the target (adj.p=0.0227). In addition, $CSDS^{SUSC}$ KO spent significantly less time in the interaction zone with the target compared to $CSDS^{RES}$ KO females (adj.p=0.004). Data are shown as mean ± SD, with *$p$<0.05, **$p$<0.01, ***p<0.001, and ****p<0.0001 indicating significance.

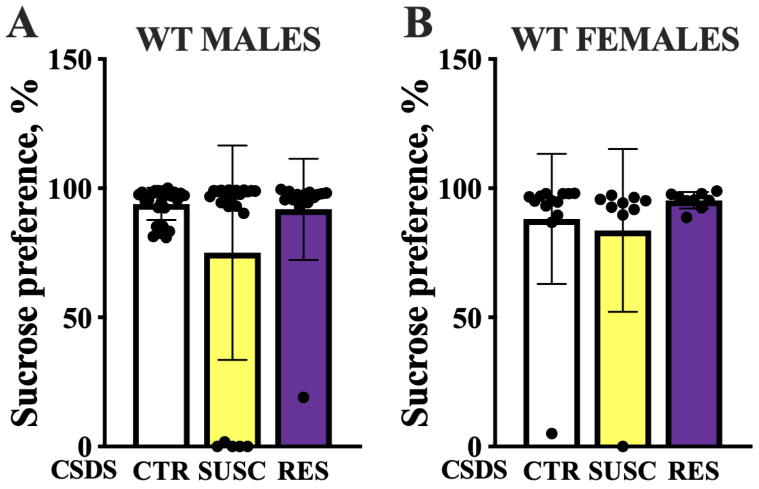


**<u>Supplementary Figure 2 -</u>** <u>Sucrose preference test.</u> No significant differences in sucrose preference were observed between $CSDS^{CTR}$, $CSDS^{SUSC}$ and $CSDS^{RES}$ groups (A – WT males, B – WT females). One-way ANOVA with Tukey's *post hoc* test. Data are shown as mean ± SD.

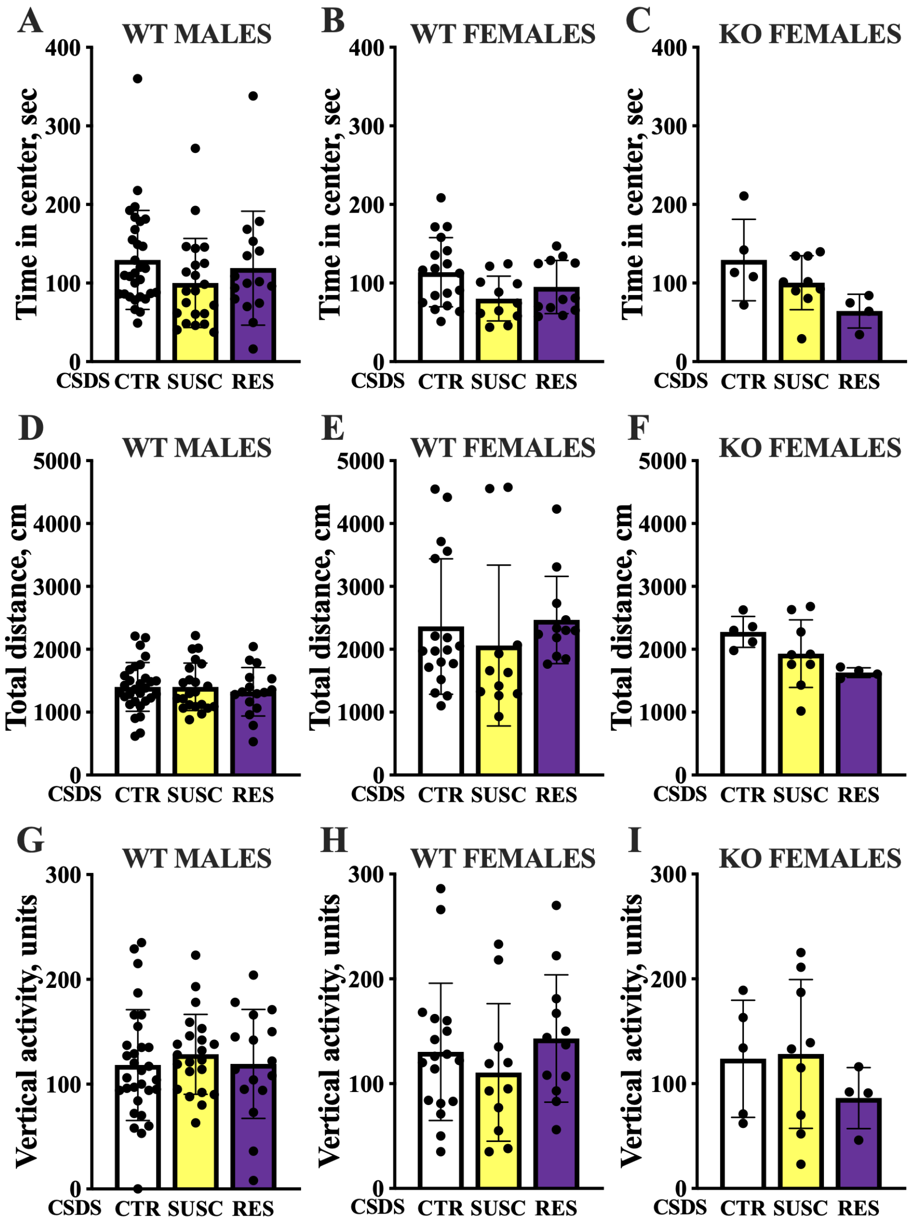


**Supplementary Figure 3 -** Open Field test (OF).

Time spent in the center zone (A – WT males, B – WT females, C – KO females) of the OF was decreased in CSDS$^{SUSC}$, but no significant differences were found between CSDS$^{CTR}$, CSDS$^{SUSC}$ and CSDS$^{RES}$ groups. The locomotor activity on the OF measured as distance moved (D – WT males, E – WT females, F – KO females) and vertical activity in the OF (G – WT males, H – WT females, I – KO females) was not significantly different among CSDS$^{CTR}$, CSDS$^{SUSC}$ and CSDS$^{RES}$ groups. One-way ANOVA with Tukey's *post hoc* test. Data are shown as mean ± SD.

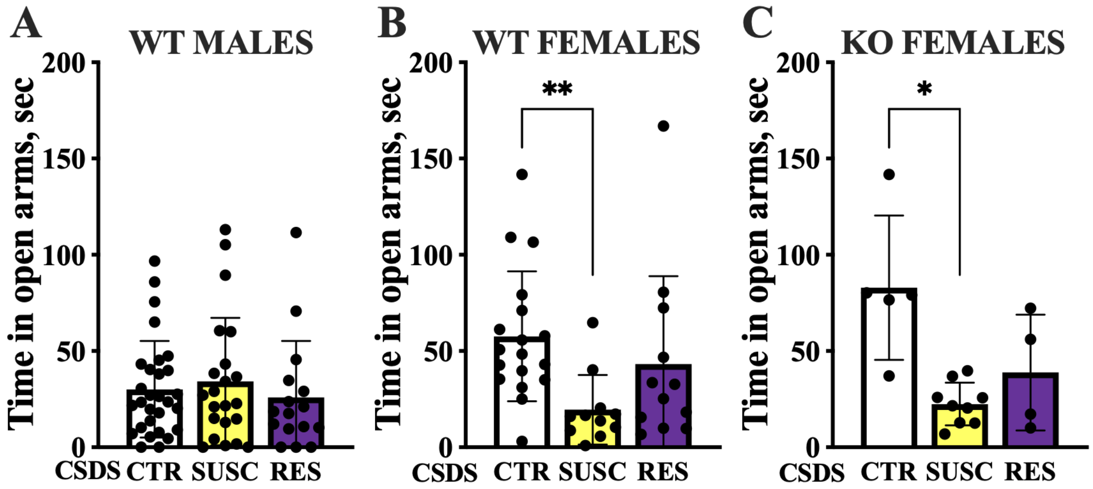


**Supplementary Figure 4 -** Elevated Plus Maze test (EPM). Time spent in the open arms of EPM was not significantly different between CSDS$^{CTR}$, CSDS$^{SUSC}$ and CSDS$^{RES}$ groups in WT males (A). In WT females (B), a Kruskal-Wallis test revealed a significant difference among groups (H(2) = 10.95, P = 0.0042), with Dunn's multiple comparisons test showing that CSDS$^{SUSC}$ WT females spent significantly less time in open arms compared to CSDS$^{CTR}$ WT females (adj.p=0.0031). In KO females (C), a similar pattern was observed, with the Kruskal-Wallis test indicating a significant difference among groups (H(2) = 8.782, P = 0.0052). *Post hoc* Dunn's test showed that that CSDS$^{SUSC}$ KO females spent significantly less time in open arms compared to CSDS$^{CTR}$ KO females (adj.p=0.0101). Data are shown as mean ± SD, with *$p$<0.05 and **$p$<0.01 indicating significance.

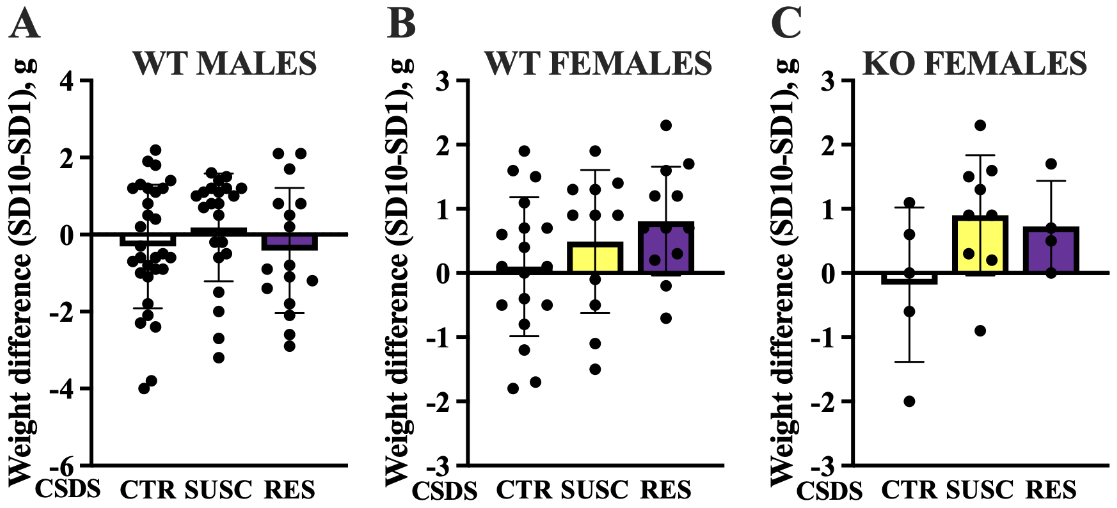


**Supplementary Figure 5 -** Weight change after CSDS. No significant differences in body weight change were observed between $CSDS^{CTR}$, $CSDS^{SUSC}$ and $CSDS^{RES}$ groups (A – WT males, B – WT females, C – KO females). However, in both WT and KO females, CSDS-defeated animals ($CSDS^{SUSC}$ and $CSDS^{RES}$) showed a trend higher weight gain compared to $CSDS^{CTR}$ between the final and initial days of social defeat. One-way ANOVA with Tukey's *post hoc* test. Data are shown as mean ± SD.

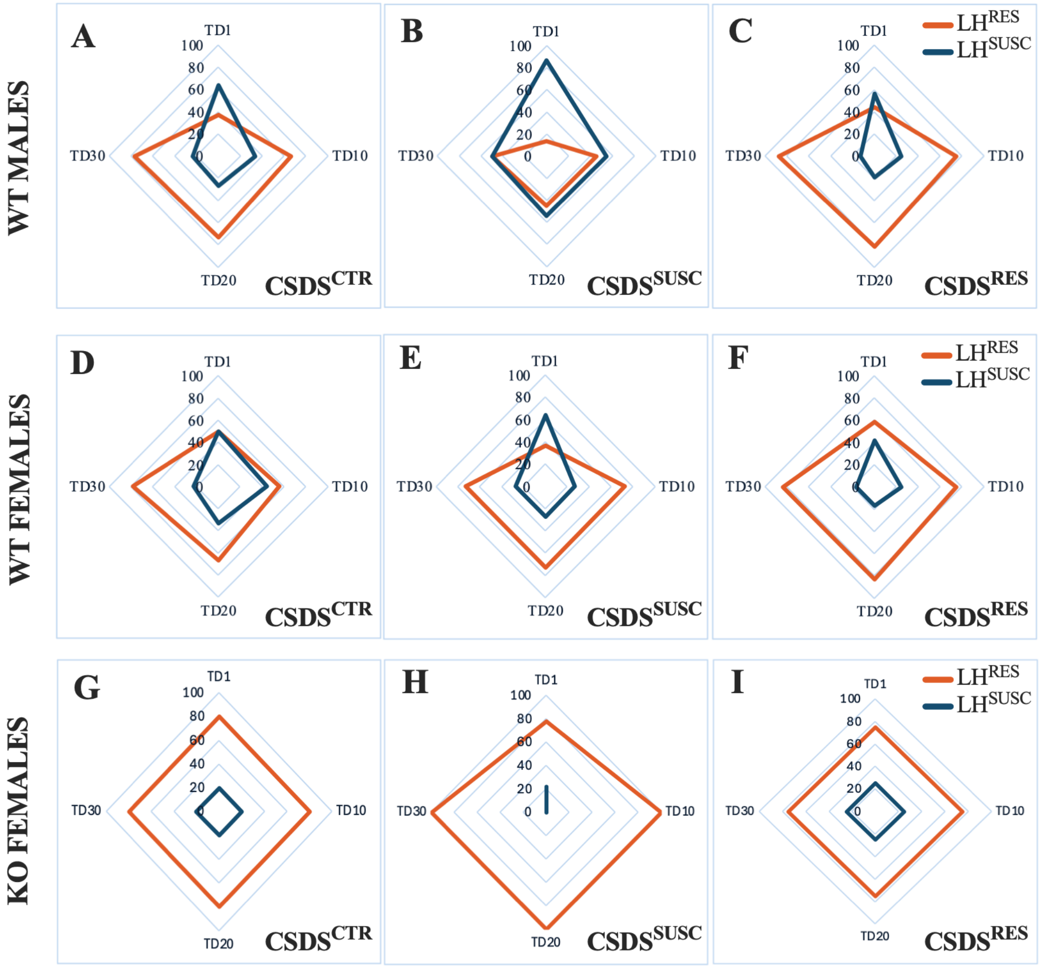


**Supplementary Figure 6 -** Radar plots representing the proportions of LH$^{RES}$ and LH$^{SUSC}$ animals among CSDS$^{CTR}$, CSDS$^{SUSC}$ and CSDS$^{RES}$ groups 1 (TD1), 10 (TD10), 20 (TD20) and 30 (TD30) days following the last LH training session (A-C – WT males, D-F – WT females, G-I – KO females).
